# Oxytocin amplifies sex differences in human mate choice

**DOI:** 10.1101/416198

**Authors:** Lei Xu, Benjamin Becker, Ruixue Luo, Xiaoxiao Zheng, Weihua Zhao, Qiong Zhang, Keith M. Kendrick

**Author notes:** Corresponding authors: Keith M. Kendrick, Address: No. 2006, Xiyuan Ave, West Hi-tech Zone, Chengdu 611731, China. Qiong Zhang Address: No. 2006, Xiyuan Ave, West Hi-tech Zone, Chengdu 611731, China.

## Abstract

Infidelity is the major cause of breakups and individuals with a history of infidelity are more likely to repeat it, but may also present a greater opportunity for short-term sexual relationships. Here in a pre-registered, double-blind study involving 160 subjects we report that while both sexes valued faithful individuals most for long-term relationships, both single men and those in a relationship were more interested in having short-term relationships with previously unfaithful individuals than women. Oxytocin administration resulted in men rating the faces of unfaithful women as more attractive but in women rating those of unfaithful men as less attractive and also finding them less memorable. Oxytocin also increased men’s interest in having short-term relationships with previously unfaithful women whereas it increased women’s interest in having long-term relationships with faithful men. Thus, oxytocin release during courtship may first act to amplify sex-dependent priorities in attraction and mate choice before subsequently promoting romantic bonds.

## 1 Introduction

Individuals who have previously been unfaithful in a relationship are over 3 times more likely to repeat this in subsequent ones (Knopp et al., 2017), and infidelity is the most common cause of divorce (Lansford, 2009). Infidelity in a partner represents a long-term relationship risk to both sexes that can particularly impact negatively on females in terms of loss of support for raising offspring but for males may also increase the risk of being cuckolded and raising another male’s offspring (Buss and Schmitt, 1993). Indeed, it is argued that this difference in the perceived risk of infidelity by the sexes is reflected in women being more concerned by emotional infidelity but men by sexual infidelity (Buss, 2018; Buss et al., 1992). However, while both sexes clearly prefer fidelity in a prospective long-term partner men are more likely to pursue short-term relationships and engage in casual sex in order to increase their reproductive potential (Buss and Schmitt, 1993; Oliver and Hyde, 1993), although women may do so to maximize their chance of reproducing with more masculine men who have the highest levels of genetic fitness (Penton-Voak et al., 1999). There is also an element of social learning in mate choice: “wanting women other men want or vice versa”, known as “mate-choice copying” (Place et al., 2010) which could be evidenced by knowledge that individuals have had multiple affairs. As Scott Fitzgerald wrote of Gatsby’s perception of Daisy in “The Great Gatsby” (Fitzgerald, 1925): “It excited him, too, that many men had already loved Daisy – it increased her value in his eyes”. Overall therefore, individuals with a previous history of infidelity could be considered as more attractive for short-term relationships, due to a greater perceived potential availability for reproduction opportunities and possibly greater genetic fitness.

In terms of the biological underpinnings of evolutionary sex-differences in human mate choice strategy, one potential candidate is the highly evolutionarily conserved neuropeptide oxytocin (OXT) which plays a key role in the formation and maintenance of affiliative and partner bonds in a number of species (Cavanaugh et al., 2014; Donaldson and Young, 2008; Kendrick et al., 2017), including humans (Preckel et al., 2014; Scheele et al., 2012, 2013), as well as in social learning (Hu et al., 2015) and conformity (De Dreu and Kret, 2016; R. Luo et al., 2017). In humans, OXT facilitates sex-dependent differences in social priorities, particularly in terms of positive or negative social attributes (Gao et al., 2016; L. Luo et al., 2017; Scheele et al., 2014). Oxytocin can also sex-dependently facilitate approach or avoidance behavior towards attractive strangers of the opposite sex although its effects can be modulated by relationship status (Scheele et al., 2012). However, it is currently unknown whether OXT may influence sex-differences in human mate-choice priorities.

Against this background we have therefore investigated whether sex-dependent biases in patterns of mate choice revealed by knowledge of previous emotional or sexual fidelity/infidelity in men and women who are either currently single or in a committed relationship, are influenced by intranasal OXT administration. We hypothesized firstly that under placebo (PLC) control conditions men would exhibit a preference for women who had been unfaithful in a previous relationship whereas women would exhibit an aversion to unfaithful men and instead prefer men who had previously been faithful. Secondly, we hypothesized that under OXT such sex differences in mate preference would be enhanced and particularly in single individuals who should have a greater priority for finding a partner than those already in an established relationship.

## 2 Methods

### 2.1 Participants

160 heterosexual human subjects (80 males, age range 18-27 years) from the University of Electronic Science and Technology of China (UESTC) were recruited to take part in a double-blind, placebo-controlled, between-subject design experiment. An initial power analysis showed that with this number of subjects the study had 80.7% statistical power for detecting treatment and sex effects with a medium effect size of 0.45 (fpower.sas). All subjects had normal or corrected-to-normal vision, were not color-blind and reported no history of or current neurological or psychiatric disorders. Subjects were free of regular and current use of medication and instructed to abstain from caffeine, nicotine and alcohol intake the day before and on the day of the experiment. None of the female subjects was pregnant or using oral contraceptives or tested at specific stages of their menstrual cycle. Using onset date of previous menses and cycle length (mean ± sem: 30.83 ± 0.37 days) provided by the subjects we estimated (backward counting (Gangestad et al., 2016)) whether they were in follicular phase (between the end of menses and ovulation, high conception risk) or luteal phase (after ovulation and before the onset of menses, low conception risk) on the experimental day (Penton-Voak et al., 1999). Eight females reported having irregular menstrual cycles and were excluded for menstrual cycle related analysis. The proportion in their follicular (*n* = 39; 22 in the OXT group) or luteal (*n* = 33; 16 in the OXT group) phases did not differ between the groups (Fisher’s exact test: *p* = 0.636, two-sided). There were no significant menstrual cycle effects found for results obtained in the study itself (see SI). Both subjects who were currently single (n = 82; 39 males) and those who were currently in a committed relationship of > 6 month duration (32.00 ± 2.45 months; n = 78; 41 males) were included since relationship status can modulate OXT effects in men (Scheele et al., 2012; Zhao et al., 2018). All single subjects were interested in finding a romantic partner and those in a relationship reported that it was a stable exclusive one (indeed subjects in a relationship scored significantly higher on the passionate love scale than single subjects (102.09 ± 1.55 vs. 96.46 ± 1.70 - *t*(158) = 2.442, *p* = 0.016, *d* = 0.387) providing further support for their being in love). All subjects signed written informed consent and received monetary compensation for their participation. The study was approved by the local ethics committee at the University of Electronic Science and Technology of China and was in accordance with the latest revision of the Declaration of Helsinki. The study was also pre-registered on the NIH registration website (Trial ID: NCT02733237; https://clinicaltrials.gov/ct2/show/NCT02733237).

To control for potential confounds, before intranasal treatment all subjects completed a range of validated questionnaires (Chinese versions) measuring mood, personality traits and attitudes toward love, trust and forgiveness (See SI). Univariate ANOVAs on questionnaires and age showed no significant differences between the OXT- and PLC-treated males and females (sex x treatment interaction: all *ps* > 0.070; See Table 1).

**Table 1.** Ages and questionnaire scores in the four experimental groups (mean±S.E.M.)

| Measurements | Placebo |  | Oxytocin |  | Sex x Treatment |
| --- | --- | --- | --- | --- | --- |
|  | Male | Female | Male | Female | *p*-value |
| Age(years) | 23.0±0.3 | 22.8±0.3 | 22.9±0.3 | 22.7±0.3 | 0.872 |
| Positive and Negative Affective Scale (PANAS) -Positive | 31.6±0.8 | 29.2±0.9 | 28.8±0.9 | 28.5±1.0 | 0.237 |
| Positive and Negative Affective Scale (PANAS) -Negative | 21.7±1.3 | 18.1±1.1 | 19.2±1.1 | 18.0±1.0 | 0.306 |
| NEO-Five Factor Inventory-Agreeableness | 42.4±0.7 | 41.4±0.6 | 40.7±0.6 | 40.9±0.8 | 0.365 |
| NEO-Five Factor Inventory-Conscientiousness | 42.7±0.8 | 41.7±0.7 | 41.4±0.8 | 42.3±0.8 | 0.220 |
| NEO-Five Factor Inventory-Extraversion | 41.2±0.8 | 38.8±1.0 | 39.9±1.0 | 40.8±0.8 | 0.070 |
| NEO-Five Factor Inventory-Neuroticism | 33.8±1.3 | 34.5±1.1 | 34.4±1.1 | 34.1±1.2 | 0.667 |
| NEO-Five Factor Inventory-Openness | 40.4±0.7 | 38.2±0.9 | 39.8±0.8 | 39.3±0.8 | 0.275 |
| Self-Esteem Scale (SES) | 30.6±0.7 | 31.3±0.6 | 30.5±0.6 | 30.8±0.8 | 0.811 |
| Interpersonal Reactivity Index (IRI) | 50.0±1.6 | 51.9±1.5 | 45.0±1.4 | 50.8±1.6 | 0.200 |
| Autism Spectrum Quotient (ASQ) | 20.0±0.7 | 20.7±0.9 | 20.7±0.6 | 19.7±0.8 | 0.254 |
| Beck Depression Inventory (BDI-II) | 8.1±0.9 | 7.9±1.2 | 8.8±1.3 | 7.2±0.9 | 0.512 |
| Liebowitz's Social Anxiety Scale (LSAS)-Avoid | 20.8±1.9 | 19.1±1.8 | 20.2±1.4 | 21.5±1.9 | 0.421 |
| Liebowitz's Social Anxiety Scale (LSAS)-Fear | 24.7±2.0 | 21.6±1.6 | 22.2±1.5 | 25.3±2.1 | 0.089 |
| Passionate Love Scale (PLS) | 103.4±2.4 | 99.0±2.3 | 97.8±2.0 | 96.7±2.5 | 0.476 |
| Love Attitude Scale (LAS)-Agape | 26.9±0.6 | 22.0±0.5 | 25.2±0.6 | 20.8±0.5 | 0.686 |
| Love Attitude Scale (LAS)-Eros | 24.1±0.5 | 23.5±0.5 | 23.4±0.6 | 23.6±0.6 | 0.529 |
| Love Attitude Scale (LAS)-Ludus | 19.3±0.7 | 19.0±0.6 | 19.9±0.5 | 18.9±0.5 | 0.573 |
| Love Attitude Scale (LAS)-Mania | 21.2±0.7 | 19.7±0.7 | 21.0±0.6 | 19.9±0.7 | 0.706 |
| Love Attitude Scale (LAS)-Pragma | 22.6±0.7 | 23.1±0.6 | 20.9±0.7 | 23.0±0.6 | 0.213 |
| Love Attitude Scale (LAS)-Storge | 22.8±0.7 | 21.5±0.7 | 21.3±0.7 | 21.7±0.8 | 0.261 |
| General Trust Scale (GTS) | 32.0±0.5 | 31.6±0.6 | 31.2±0.6 | 31.8±0.7 | 0.380 |
| Tendency to Forgive Scale (TTF) | 14.3±0.5 | 14.3±0.5 | 14.1±0.6 | 13.9±0.7 | 0.846 |
| Attitudes toward Forgiveness Scale (ATF) | 28.5±0.7 | 27.9±0.7 | 27.4±0.6 | 26.5±0.7 | 0.808 |
| Trait Forgivingness Scale (TFS) | 32.7±0.9 | 32.5±0.9 | 32.0±0.8 | 30.9±1.0 | 0.614 |

### 2.2 Intranasal administration

Subjects were randomly assigned to receive intranasal administration of either OXT (*n* = 80, 40 males and 40 females; 40 IU; OXT-Spray, Sichuan Meike Pharmaceutical Co. Ltd, China; 5 puffs of 4 IU per nostril with a 30s interval between each puff) or PLC (*n* = 80, 40 males and 40 females; identical sprays with the same ingredients other than the neuropeptide, i.e., glycerin and sodium chloride) following a standardized protocol (Guastella et al., 2013). In previous studies, we have found similar behavioral and neural effects of 24 and 40 IU OXT doses, although the higher dose tended to produce more consistent results (Geng et al., 2018; Xu et al., 2015; Zhao et al., 2017) and this was recently supported by a study from another group showing dose-dependent effects using these same doses (Shin et al., 2018). We therefore decided to use the higher 40 IU dose here to try and maximize effects. Although we could not measure blood or cerebrospinal fluid OXT concentrations following intranasal application other studies have reported that they produce only relative small increases within the general physiological range (Quintana et al., 2018; Striepens et al., 2013). While it is currently unclear whether intranasal OXT produces direct effects on the brain or also indirectly via peripheral effects, it has been established that OXT administered via this route does enter into the brain cerebroventricular system in monkeys (Lee et al., 2018) and alters cerebral blood flow in an extensive number of brain regions known to express OXT receptor mRNA in humans (Paloyelis et al., 2016). A recent study comparing functional and brain effects of intranasal and intravenous OXT administration also only found effects when it is given intranasally (Quintana et al., 2016). For allocation of the participants to the two treatment groups a computer-generated list of random numbers was used (groups, n = 2; numbers per group, n = 40; simple randomization). Treatment allocation was done by an experimenter not involved in data acquisition and analyses. Subjects and experimenter were blind to drug condition. In post experiment interviews subjects were unable to guess better than chance whether they had received OXT or PLC treatment (81 subjects guessed correctly; χ^2^ = 0.025, p = 0.874). In line with standardized recommendations (Guastella et al., 2013) and two studies reporting pharmacodynamics of central effects of intranasal OXT in humans (Paloyelis et al., 2016; Spengler et al., 2017) the experimental paradigm started 45 minutes after intranasal treatment.

### 2.3 Stimuli

Before the formal experiment, we generated 54 sentences describing a behavior indicative of fidelity or infidelity (either emotional or sexual; 12~14 sentences for each behavior type) that a male or female individual had performed during a past relationship. Sexual and emotional infidelity were defined as in Takahashi et al (Takahashi et al., 2006). Sexual infidelity (or fidelity) included situations where a (or no) sexual relationship or deep physical contact with other members of the opposite sex was indicated explicitly or implicitly. Emotional infidelity (or fidelity) included situations indicating some (or no) form of romantic emotional response or commitment to other members of the opposite sex. Each sentence was written in Chinese, used the past tense and had male and female versions (i.e. “She…….” for male subjects in the study and “He……” for female subjects). In a pre-study, an independent sample of forty volunteers (21 males) were asked to decide whether the behavior described was an example of emotional or sexual infidelity/fidelity and also to rate how strong it was using a 9-point scale. Based on the data from this pre-study, we selected 40 sentences (10 for each behavior type) with a high discrimination between sexual and emotional fidelity or infidelity (i.e. all the chosen sentences were correctly classified as representing fidelity or infidelity behaviors by the raters and with a mean accuracy of 87.6% for distinguishing emotional from sexual examples). There were no differences between male and female examples in terms of discrimination accuracy or strength (all *ps* > 0.258). Table S1 gives examples of the emotional or sexual fidelity/infidelity behavior sentences.

Facial images of 80 males and 80 females with neutral expressions were selected from an in-house database of 260 face images following a pilot rating by 36 subjects (17 males) of valence, attractiveness, likeability, trustworthiness of the faces from the opposite sex as well as how aroused they were by them. All face images were carefully edited (removing accessories or background details, but keeping hair, ears and neck) and presented in full color at a 600×800 Pixel resolution on a black background (faces life-size). All selected faces were rated as having a neutral valence (range 4.3-6.0; mean = 5.09) and average attractiveness (range 4.0-5.9, mean= 4.79), likeability (range 4.1-5.8, mean= 4.81) and trustworthiness (range 4.2-6.0, mean= 5.07). Half of the faces used for the rating task were divided randomly into four groups (i.e. 10 faces per group for each sex). Mean valence, attractiveness, likeability, trustworthiness and arousal ratings of the faces in each group did not differ significantly for both male and female faces (ANOVAs all *ps* > 0.964). Each group of faces was assigned for pairing with sentences describing one of the four different fidelity/infidelity types. Additionally, to control for possible face/sentence-group differences, the pairings of face group and sentence type were randomized across individual subjects in the main study. The remaining faces were used as novel stimuli in the recognition memory test and had equivalent valence, attractiveness, likeability, trustworthiness and arousal ratings compared to the faces paired with sentences for both sexes (all *ps* > 0.661).

### 2.4 Procedure

The experimental task (see Fig. 1) was presented on a computer with a 27-inch monitor (screen resolution: 1920*1080 pixels; refresh rate: 60 Hz). In the rating task, subjects viewed neutral expression face pictures of 40 unfamiliar members of the opposite sex with average attractiveness paired with verbal information describing examples of how they had been either emotionally or sexually faithful or unfaithful during a previous relationship (see Table S2). We included fidelity type as a factor since previous research has reported that men are more influenced by sexual infidelity and women by emotional infidelity (Buss, 2018; Buss et al., 1992). Subjects were told that these individuals were currently single and instructed to view their faces, read the sentences describing their previous behavior silently and then rate (on a 9-point scale) their facial attractiveness, likeability, trustworthiness and arousal elicited by them based on their overall impression of them. Next, subjects were asked whether they would like to have a short- or long-term romantic relationship with the person (response options: “yes”, “maybe” or “no” - see Fig.1). There was no time limitation for subjects’ responses. The percentage of “yes/maybe” responses made by each subject for each condition indicated their willingness to have a relationship with this kind of person.

**Fig. 1.**
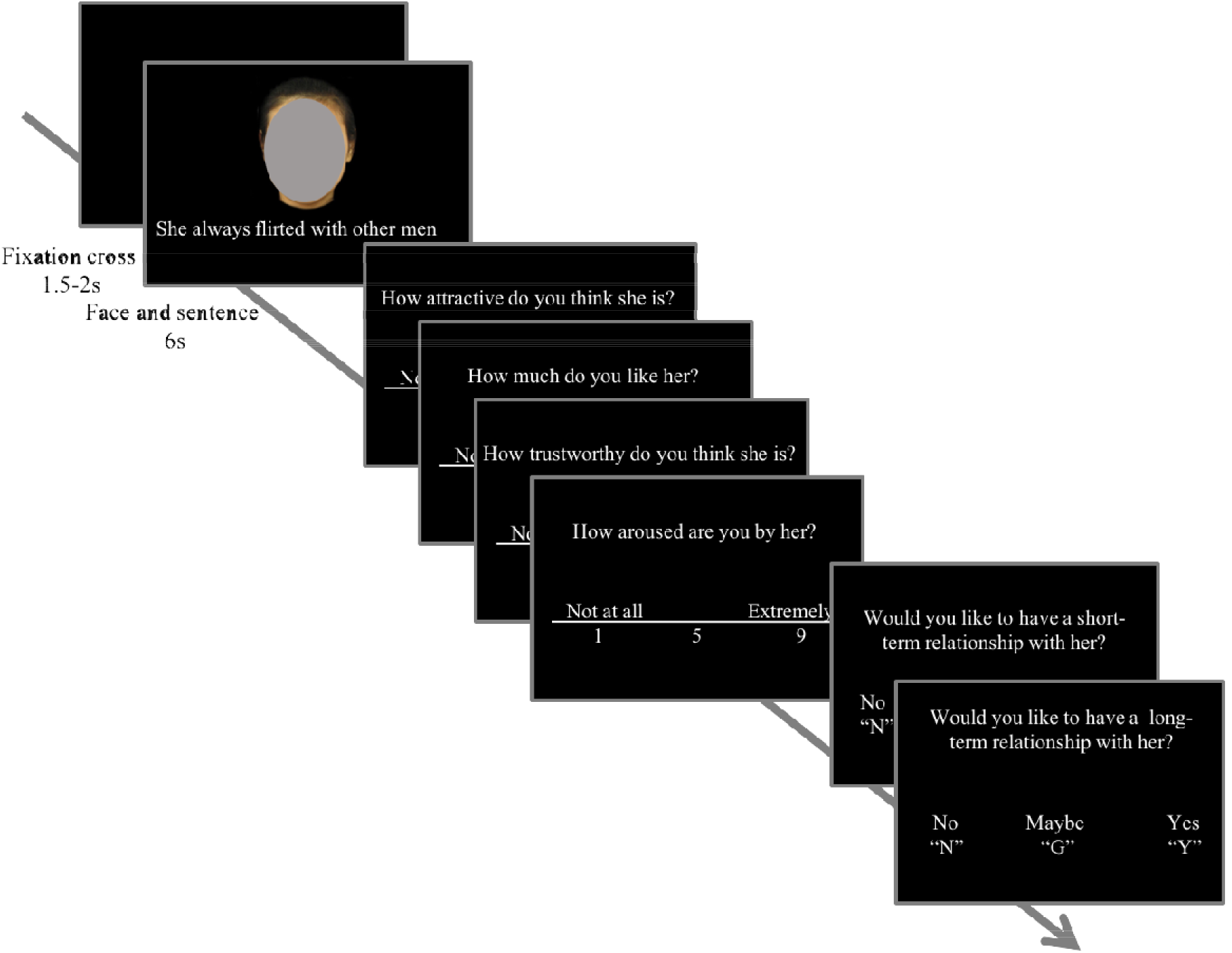
Example of a single trial in the rating task. Following a 1.5~2 second fixation cross, each facial picture (unknown, opposite sex) was shown for 6 seconds and paired with a sentence describing a behavior indicative of fidelity or infidelity (either emotional or sexual) he/she exhibited during a previous relationship. Each subject viewed 10 trials for each fidelity/infidelity type - emotional fidelity, sexual fidelity, emotional infidelity and sexual infidelity.

Finally, subjects completed a surprise recognition memory test for these 40 faces intermixed with another 40 novel faces (order of stimuli randomized). Each trial started with a 600-800 ms fixation cross followed by a face presented for 1500 ms and subjects responded whether the face was familiar or not without any time limitation. Four subjects had to be excluded from this part of the analysis due to technical failures during data acquisition.

### 2.5 Statistical Analysis

All data analyses were performed using SPSS 23.0 software (SPSS Inc., Chicago, Illinois, USA). In all cases, data from rating scores, recognition memory accuracy and percentage of “yes/maybe” responses for having a short- or long-term relationship with a target individual were subjected to four (analysis of the PLC group alone) or five (analysis of the PLC vs. OXT treatment groups) factor repeated-measures ANOVAs and significant (*p* < 0.05) main effects and relevant interactions were reported. Significant interactions were explored using Simple Effect Tests, which were all Bonferroni-corrected for multiple comparisons. For both ANOVAs and post-hoc tests measures of effect size are given (Partial eta squared (*η*^2^_p_) or Cohen’s *d*). Small, medium, and large effects were represented respectively as 0.01, 0.06, and 0.14 for *η*^2^_p_, 0.20, 0.50, and 0.80 for Cohen’s *d* (Cohen, 1988).

## 3 Results

### 3.1 Sex-differences on the impact of knowledge of previous fidelity or infidelity

To identify treatment-independent sex differences on evaluations of a potential partner who had previously displayed emotional or sexual fidelity or infidelity in a relationship, we first analyzed data from the PLC control group using four-way repeated-measures ANOVAs with fidelity (fidelity vs. infidelity) and type (emotional vs. sexual) as within-subject factors and sex and relationship status as between-subject factors.

For attractiveness, likeability, trustworthiness and arousal ratings of the face pictures paired with examples of fidelity or infidelity behaviors there were no significant fidelity x sex interactions (all *ps* > 0.328). However, there was a significant type x sex interaction for likeability ratings (*F*(1,76) = 5.447, *p* = 0.022, *η*^2^_p_ = 0.067). Post hoc comparisons revealed that women rated men who showed emotional fidelity or infidelity (4.15 ± 0.10, 95% CI = [3.96, 4.34]) more likeable than those who showed sexual fidelity or infidelity (3.94 ± 0.10, 95% CI = [3.74, 4.15]; *p* = 0.001, *d* = 0.366 – see Fig. S1a). Thus, the most likeable potential partners for men were those who showed sexual fidelity while for women they were those who showed emotional fidelity. There was also a similar trend for this with attractiveness ratings although the interaction was only marginally significant (*F*(1,76) = 3.731, *p* = 0.057, *η*^2^_p_ = 0.047 – see Fig. S1b). There were no significant interactions involving type and sex for recognition memory accuracy (all *ps* > 0.094).

For short-term relationship preferences, analysis in the PLC group revealed a significant fidelity x sex interaction (*F*(1,76) = 8.807, *p* = 0.004, *η*^2^_p_ = 0.104). Post-hoc Bonferroni corrected comparisons showed that 31.8 ± 3.6% (95% CI = [24.6%, 38.9%]) of responses made by men expressed interest (i.e. “yes” or “maybe” decisions) in having a short-term relationship with an unfaithful individual, whereas only 17.0 ± 3.6% (95% CI = [9.8%, 24.1%]) of responses made by women did (*p* = 0.005, *d* = 0.658 – see Fig. 2). There were no sex-differences for long-term relationship preferences (all *ps* > 0.109), with both men (44.1 ± 4.9%, 95% CI = [34.4%, 53.9%]) and women (47.9 ± 4.9%, 95% CI = [38.1%, 57.6%]) showing an equivalent and greater preference for partners exhibiting previous fidelity (*p* = 0.589).

**Fig. 2.**
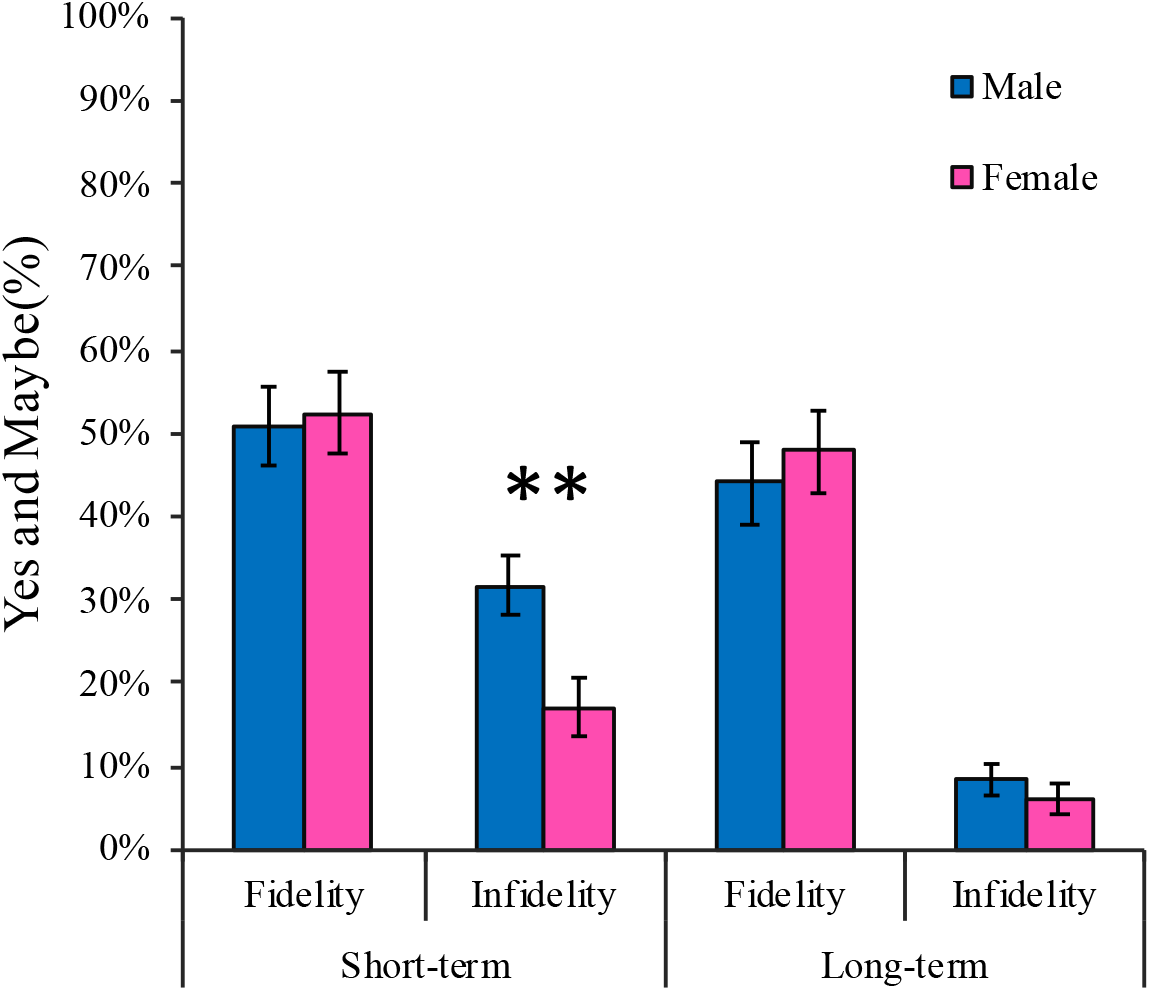
Sex difference in the percentage of yes/maybe responses made by subjects for having a short-term, but not long-term, relationship with individuals showing previous infidelity in the placebo treated group. Data from single individuals and those in a relationship are combined. Bars represent means and standard errors. \*\**p* < 0.01 for males vs. females.

A separate analysis on female subjects found no evidence for a significant influence of menstrual cycle stage (i.e. whether women were at a stage representing either a high or low risk of conception) on any of the measures taken (see SI).

### 3.2 Effects of intranasal oxytocin on sex-differences in mate choice

To examine the effects of OXT on evaluations of potential partners showing previous fidelity or infidelity, five way repeated-measures ANOVAs with fidelity (fidelity vs. infidelity) and type (emotional vs. sexual) as within-subject factors and treatment, sex and relationship status as between-subject factors were performed on rating scores, recognition memory accuracy and percentage of “yes/maybe” responses for having a short- or long-term relationship with a target individual.

There were significant fidelity x treatment x sex interactions for attractiveness (*F*(1,152) = 8.454, *p* = 0.004, *η*^2^_p_ = 0.053 – see Fig. 3a) and likeability ratings (*F*(1,152) = 6.694, *p* = 0.011, *η*^2^_p_ = 0.042 – see Fig. 3b). Post-hoc Bonferroni corrected comparisons showed that in men OXT increased both face attractiveness (*p* = 0.047, *d* = 0.421) and likeability (*p* = 0.017, *d* = 0.513) of previously unfaithful women, while in women OXT decreased face attractiveness (*p* = 0.016, *d* = 0.529) and likeability (*p* = 0.181) of previously unfaithful men. Thus, unlike the PLC group, in the group treated with OXT there were significant sex differences in face attractiveness (*p* < 0.001, *d* = 1.033) and likeability (*p* < 0.001, *d* = 1.006) of previously unfaithful individuals. There were no significant OXT effects on face attractiveness or likeability ratings given to previously faithful men and women (all *ps* > 0.461). And OXT did not alter the pattern of female subjects giving higher attractiveness or likeability ratings than men for emotionally compared to sexually faithful individuals (interactions involving type, treatment and sex: all *ps* > 0.260). No significant interaction effects involving treatment, sex, fidelity or type were found for trustworthiness (all *ps* > 0.075) or arousal ratings (all *ps* > 0.134) indicating that sex-dependent effects of OXT on attraction ratings were specific.

**Fig. 3.**
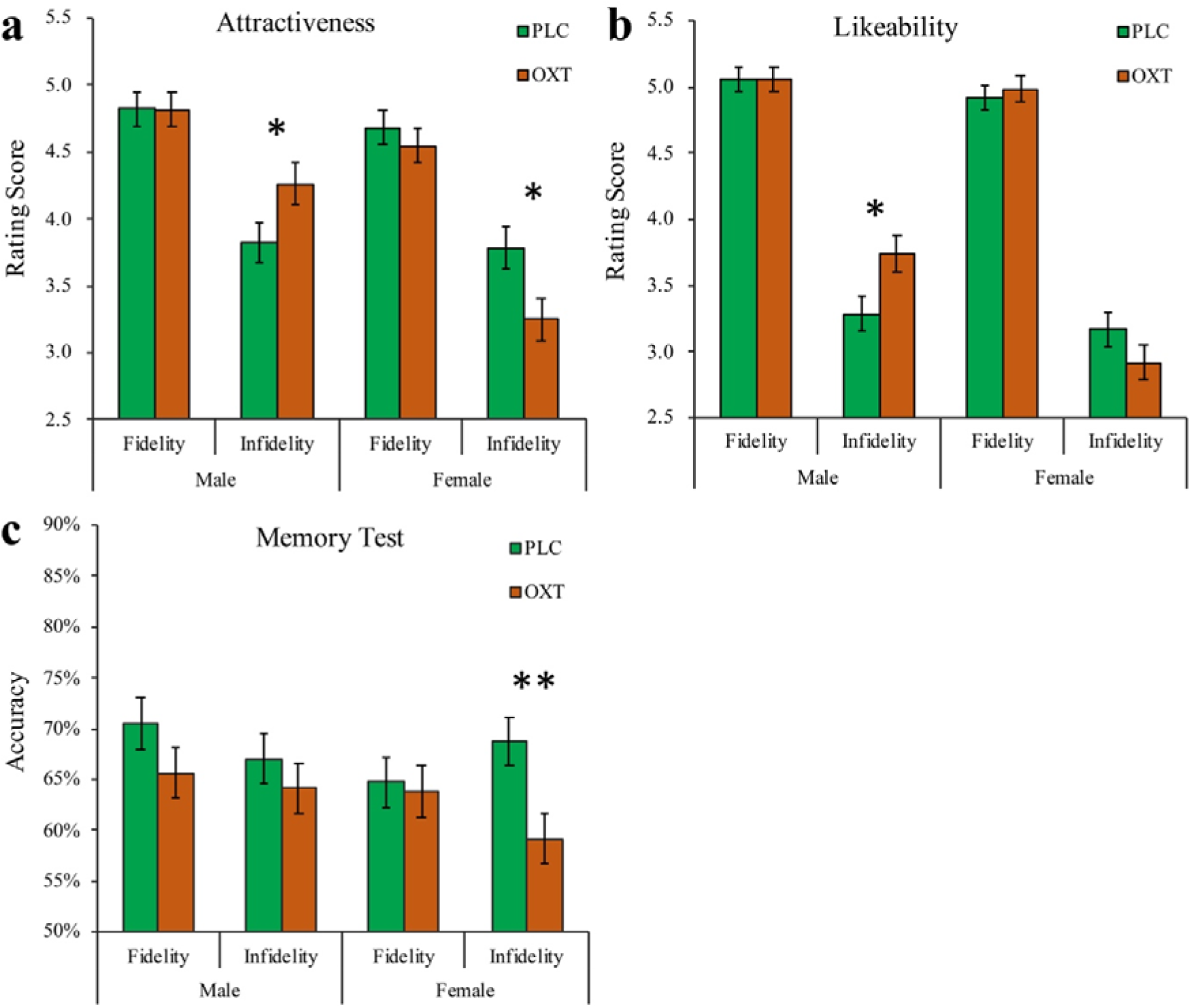
Effects of oxytocin (OXT) on attractiveness (**a**) likeability **(b)** and recognition memory (**c**) for faces of the opposite sex associated with previous fidelity or infidelity in all male and female subjects. Bars represent means and standard errors. \**p* < 0.05, \*\**p* < 0.01 OXT vs. placebo (PLC).

Analysis of recognition memory accuracy for faces revealed a significant fidelity x treatment x sex interaction (*F*(1,148) = 6.036, *p* = 0.015, *η*^2^_p_ = 0.039; note: for this analysis 4 subjects were excluded due to incomplete data). Post-hoc Bonferroni corrected comparisons demonstrated that women in the OXT group (59.1 ± 2.5%, 95% CI = [54.3%, 64.0%]) were less likely than women in the PLC group (68.8 ± 2.4%, 95% CI = [63.9%, 73.6%]) to remember the faces of individuals who had previously exhibited infidelity (*p* = 0.006, *d* = 0.608 – see Fig. 3c). Oxytocin therefore effectively increased the chances that women would only remember men with a history of being faithful. No other significant interaction effects involving treatment, sex, fidelity or type were found (all *ps* > 0.243).

For short-term relationship preference, analysis revealed a fidelity x treatment x sex x relationship status interaction (*F*(1,152) = 4.082, *p* = 0.045, *η*^2^_p_ = 0.026). Post-hoc Bonferroni corrected comparisons showed that the percentage of yes/maybe responses given by single men for having a short-term relationship with unfaithful women were increased from 30.0 ± 4.9% (95% CI = [20.4%, 39.6%]) in the PLC group to 45.8 ± 5.0% (95% CI = [35.9%, 55.7%]) in the OXT group (*p* = 0.025, *d* = 0.643 – see Fig. 4a).No other significant interaction effects involving treatment, sex, fidelity or type were found (all *ps* > 0.06).

**Fig. 4.**
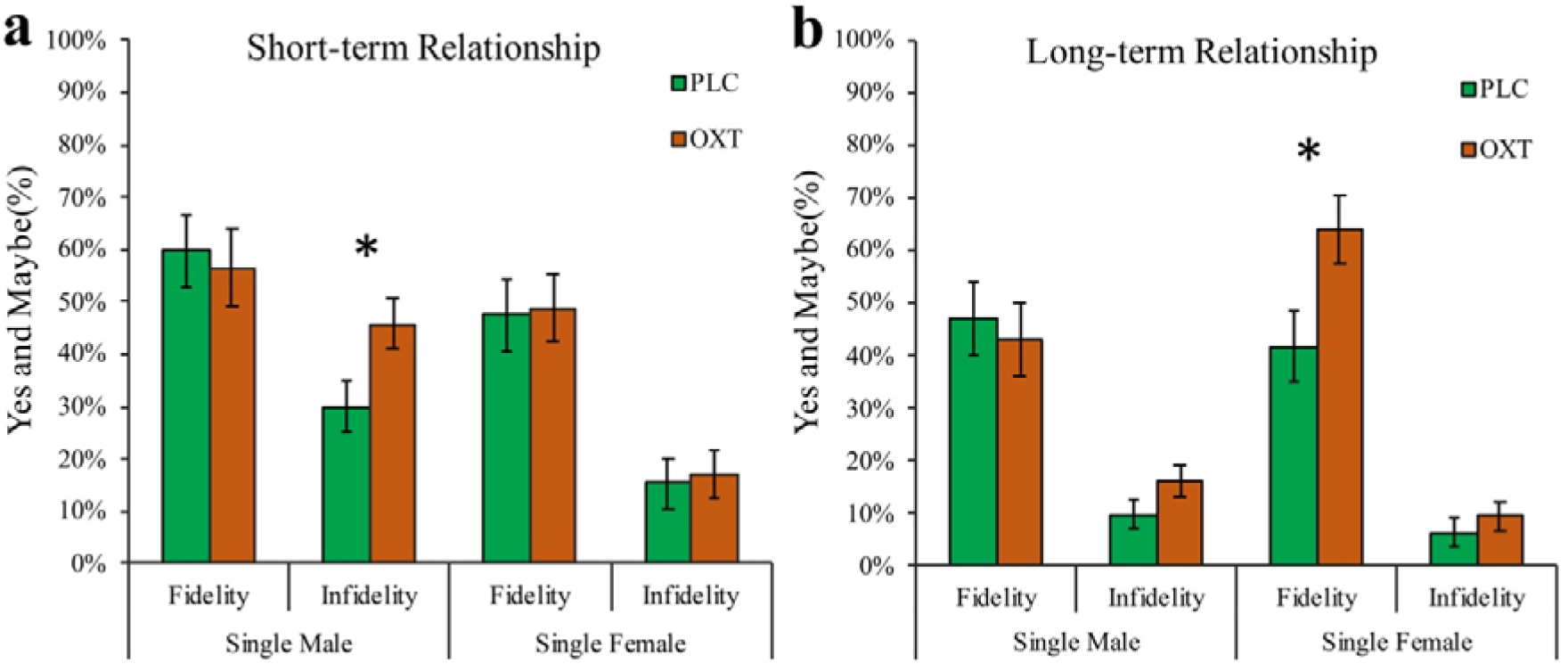
Effect of oxytocin (OXT) on percentage of yes/maybe responses in single male and female subjects for having a short-term (**a**) or long-term relationship (**b**) with an individual of the opposite sex associated with previous fidelity or infidelity. Bars represent means and standard errors. \**p* < 0.05 OXT vs. placebo (PLC).

For interest in having a long-term relationship there was a significant fidelity x treatment x sex x relationship status interaction (*F*(1,152) = 5.439, *p* = 0.021, *η*^2^_p_ = 0.035). Post-hoc Bonferroni corrected comparisons showed that the percentage of yes/maybe responses made by single women for having a long-term relationship with faithful men were increased from 41.8 ± 6.9% (95% CI = [28.2%, 55.3%]) in the PLC group to 64.1 ± 6.4% (95% CI = [51.5%, 76.8%]) in the OXT group (*p* = 0.018, *d* = 0.699 – see Fig. 4b). No other significant interaction effects involving treatment, sex, fidelity or type were found (all *ps* > 0.266).

A separate analysis on female subjects found menstrual cycle stage did not influence the OXT effects found above (see SI).

## 4 Discussion

Overall, our findings demonstrate firstly that knowledge of previous fidelity and infidelity in a prospective heterosexual partner effectively reveals sex differences in mate choice strategy. Thus, men in the control PLC treated group generally exhibited greater interest in having a short-term relationship with previously unfaithful individuals than women, and independent of relationship status. There was no sex difference in the context of long-term relationships, with both sexes showing an equivalent and greater preference for partners exhibiting previous fidelity. Following OXT administration both men and women respectively exhibited enhanced and reduced attraction to unfaithful individuals and women also found them less memorable. Additionally, single men showed an increased preference for have a short-term relationship with previously unfaithful women whereas single women showed an increased preference for having a long-term relationship with previously faithful men.

In support of our hypothesis our findings in the PLC group demonstrate a clear sex-dependent bias in mate choice with men expressing a greater interest than women in having short-term relationships with previously unfaithful individuals. This therefore tends to support proposed evolutionary sex-differences in mate choice priorities (Buss & Schmitt, 1993; Oliver & Hyde, 1993). In addition, we found some evidence to support previous studies arguing that a sex difference in the perceived risk of infidelity is reflected in women being more concerned by emotional infidelity but men by sexual infidelity (Buss, 2018; Buss et al., 1992). Females more liked emotionally faithful males and males more liked sexually faithful females.

Again in support of our original hypothesis OXT administration increased sex-differences in mate-choice priorities. Thus, in contrast to the PLC group, subjects in the OXT group exhibited sex-differences in the influence that knowledge of previous fidelity or infidelity had on attractiveness and likeability ratings and memory for prospective partners. Importantly however, OXT administration had no effect on potential confounders such as arousal and trustworthiness ratings and effects were also independent of relationship status. More specifically, OXT increased men’s attractiveness and likeability ratings of previously unfaithful women but correspondingly decreased those for unfaithful men by women. Furthermore, following OXT administration women found the face pictures of men associated with previous infidelity less memorable, suggesting that they would be more likely to only remember faithful individuals. Interestingly however, OXT did not alter the sex-specific preferences for the attractiveness and likeability ratings given to individuals who had previously exhibited emotional (female) as opposed to sexual (male) fidelity. This may reflect the fact that the sex-dependent effects of OXT were mainly in the context of interest in previous infidelity or that it may have less influence on such strongly established within-sex patterns of preference. Both the sex-differences observed in the PLC group and in response to OXT treatment were robust with all achieving medium or large effect sizes, thereby confirming the appropriateness of the power analysis for the study.

While the sex-dependent effects of OXT on attraction and likeability ratings and memory for faces occurred irrespective of relationship status, those for increasing interest in having short or long-term relationships were restricted to single individuals. This finding supports our hypothesis that OXT would enhance sex-dependent social and reproductive priorities (Gao et al., 2016; Hurlemann and Scheele, 2016) but particularly in single individuals who have a higher priority for seeking a potential partner than those already in an established relationship. That OXT primarily increased single men’s interest in having short-term relationships with women who had previously been unfaithful may reflect a high priority for gaining sexual access to females. Similarly, single women’s increased interest in faithful males, and decreased interest in and memory for unfaithful ones, may reflect both a higher priority for avoiding potential philandering males and preference for faithful individuals who are more likely to provide a stable and secure relationship.

Oxytocin release associated with partner bonding across species is primarily evoked by mating or sexual arousal as well as by affective touch (Hurlemann and Scheele, 2016; Li et al., 2019), and can even occur in response to visual cues from the face (Fabre-Nys et al., 1997). While there is some evidence that OXT can increase the perceived attractiveness of the faces of unfamiliar members of the opposite sex (Hurlemann and Scheele, 2016) our current findings emphasize that its release during initial social interactions might serve to focus attention on pertinent information concerning a prospective partner’s behavior and history and not merely on their physical appearance. Indeed, previous studies have also demonstrated that intranasal OXT administration can potently, and sex-dependently, alter behavioral and neural responses to faces when they are paired with information on positive or negative social qualities (Gao et al., 2016) and reduce recognition speed for positive romantic and bonding-related words (Unkelbach et al., 2008). Thus, while OXT release can ultimately promote the formation of partner bonds, it may first play a key role in highlighting the attractiveness of personal characteristics in a prospective partner which best match an individual’s current priorities.

The current study has several potential limitations. Firstly, the paradigm used of rating attractiveness and mating preference for individuals based purely on their face pictures associated with a verbal descriptor is commonly used, it is relatively artificial and it is possible that results in contexts involving real social interactions might have been different. Secondly, the subjects used in the study were primarily students and those in a relationship had relatively short durations (>6 months). It is possible that both sex-differences in the PLC group and the effects of OXT might have been influenced by age and also relationship durations.

## 5 Conclusions

In summary, our findings demonstrate firstly that in the context of knowledge of a prospective partner’s fidelity or infidelity in previous relationships men who are either single or in a relationship are more interested than women in having short-term relationships with unfaithful individuals. Following OXT treatment this sex difference is both amplified and extended such that men are even more interested in having short-term relationships with unfaithful individuals and women even less so. Furthermore, women exhibit a greater interest in faithful individuals than do men in the context of long-term relationships. In terms of mate choice therefore, the sexes exhibit a differential interest in prospective partners who are “stayers” or “strayers”, and for single individuals with a current priority for finding a prospective partner OXT release during romantic encounters may act to further widen this sex difference. Thus, OXT release may function first to influence sex-dependent mate-choice priorities before subsequently promoting romantic bonds with the most appropriate partners.

## Acknowledgements

We thank Professor Trevor Robbins for valuable discussions and suggestions on the paper and its findings. This project was supported by National Natural Science Foundation of Science (NSFC) grant number 31530032.

## Author Contributions

LX and KMK designed the experiment. LX, RL, XZ and WZ carried out the experiment. LX, KMK, BB and QZ analyzed the experiment and LX, KMK and BB wrote the paper. All authors contributed to the conception of the study and approved the paper.

## Supporting Information

### Supplementary Methods

To control for potential confounds, before intranasal treatment all subjects completed a range of validated questionnaires (Chinese versions) measuring mood, personality traits and attitudes toward love, trust and forgiveness. These included: Positive and Negative Affective Schedule – PANAS (Watson et al., 1988); NEO-Five Factor Inventory – NEO-FFI (Costa and Mccrae, 1989); Self-Esteem Scale – SES (Rosenberg, 1965); Interpersonal Reactivity Index – IRI (Davis, 1980); Autism Spectrum Quotient – ASQ (Baron-Cohen et al., 2001); Beck’s Depression Inventory – BDI (Beck et al., 1996); Leibowitz’s Social Anxiety Scale – LSAS (Liebowitz, 1987); Passionate Love Scale – PLS (Hatfiled and Sprecher, 1986); Love Attitude Scale – LAS (Hendrick and Hendrick, 1986); General Trust Scale – GTS (Siegrist et al., 2005); Tendency to Forgive Scale – TTF (Brown, 2003); Attitudes toward Forgiveness Scale – ATF (Brown, 2003); Trait Forgivingness Scale – TFS (Berry et al., 2005). Univariate ANOVAs on questionnaires and age showed no significant differences between the OXT- and PLC-treated males and females (sex x treatment interaction: all *ps* > 0.070; See Table 1).

### Supplementary Results

Repeated-measures ANOVAs added menstrual cycle as a between-subject factor in female subjects suggested that the stage of their menstrual cycle did not influence our findings. There were no significant interactions of menstrual cycle and fidelity or interactions of menstrual cycle and type for rating scores, memory and short-term or long-term relationship preferences in the PLC group (all *ps* > 0.100). For the effects of OXT there were also no significant interactions of menstrual cycle, treatment and fidelity for rating scores, memory and short-term or long-term relationship preferences (all *ps* > 0.390).

---

**Table S1.**
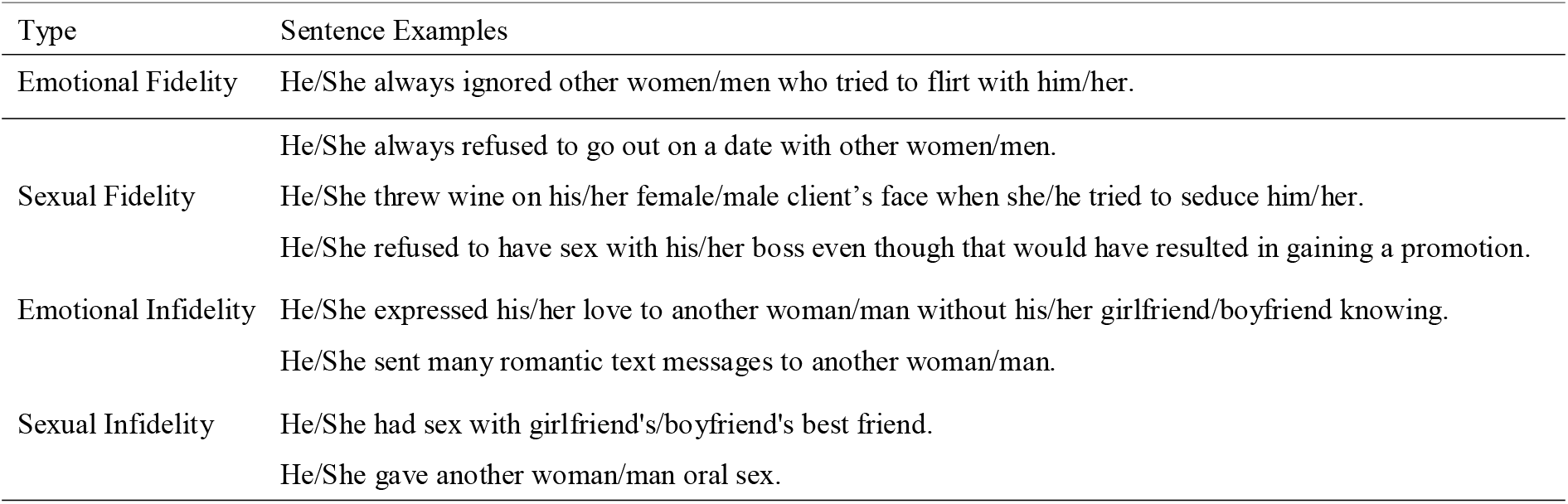
Examples of sentences describing sexual and emotional fidelity or infidelity

**Fig. S1.**
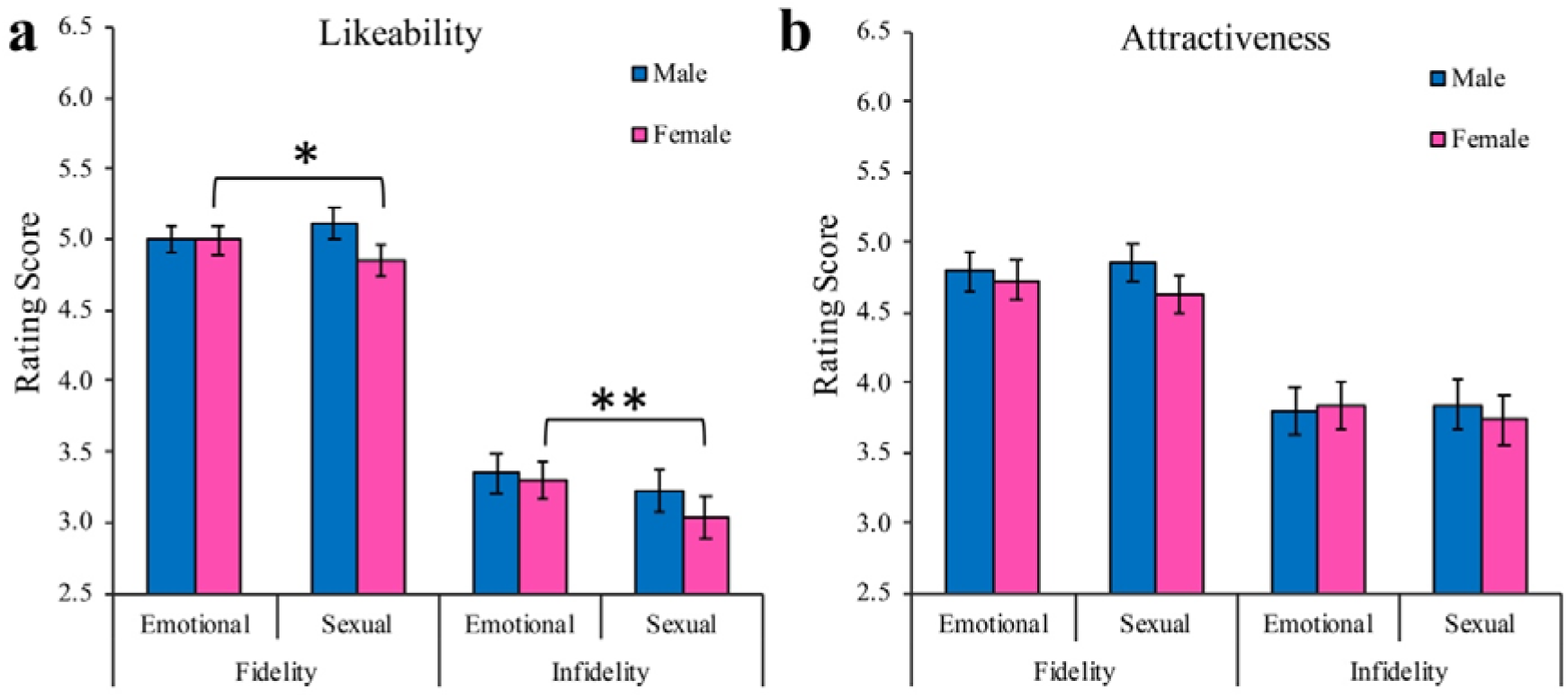
Sex difference in likeability and attractiveness ratings in the placebo (PLC) treated group. \**p* < 0.05, \*\**p* < 0.01.

## References

Buss, D.M., 2018. Sexual and emotional infidelity: Evolved gender differences in jealousy prove robust and replicable. Perspect. Psychol. Sci. 13, 155–160. https://doi.org/10.1177/1745691617698225

Buss, D.M., Larsen, R.J., Westen, D., Semmelroth, J., 1992. Sex differences in jealousy: Evolution, physiology, and psychology. Psychol. Sci. 3, 251–256. https://doi.org/10.1111/j.1467-9280.1992.tb00038.x

Buss, D.M., Schmitt, D.P., 1993. Sexual Strategies Theory: An evolutionary perspective on human mating. Psychol. Rev. 100, 204–232. https://doi.org/10.1037/0033-295X.100.2.204

Cavanaugh, J., Mustoe, A.C., Taylor, J.H., French, J.A., 2014. Oxytocin facilitates fidelity in well-established marmoset pairs by reducing sociosexual behavior toward opposite-sex strangers. Psychoneuroendocrinology 49, 1–10. https://doi.org/10.1016/j.psyneuen.2014.06.020

Cohen, J., 1988. The Effect Size index: d, in: Statistical Power Analysis for the Behavioral Sciences. pp. 20–26.

De Dreu, C.K.W., Kret, M.E., 2016. Oxytocin conditions intergroup relations through upregulated in-group empathy, cooperation, conformity, and defense. Biol. Psychiatry 79, 165–173. https://doi.org/10.1016/j.biopsych.2015.03.020

Donaldson, Z.R., Young, L.J., 2008. Oxytocin, Vasopressin, and the Neurogenetics of Sociality. Science. 322, 900–904. https://doi.org/10.1126/science.1158668

Fabre-Nys, C., Ohkura, S., Kendrick, K.M., 1997. Male faces and odours evoke differential patterns of neurochemical release in the mediobasal hypothalamus of the ewe during oestrus: an insight into sexual motivation? Eur. J. Neurosci. 9, 1666–1677. https://doi.org/10.1111/j.1460-9568.1997.tb01524.x

Fitzgerald, F.S., 1925. The Great Gatsby. Charles Scribner’s Sons, New York.

Gangestad, S.W., Haselton, M.G., Welling, L.L.M., Gildersleeve, K., Pillsworth, E.G., Burriss, R.P., Larson, C.M., Puts, D.A., 2016. How valid are assessments of conception probability in ovulatory cycle research? Evaluations, recommendations, and theoretical implications. Evol. Hum. Behav. 37, 85–96. https://doi.org/10.1016/j.evolhumbehav.2015.09.001

Gao, S., Becker, B., Luo, L., Geng, Y., Zhao, W., Yin, Y., Hu, J., Gao, Z., Gong, Q., Hurlemann, R., Yao, D., Kendrick, K.M., 2016. Oxytocin, the peptide that bonds the sexes also divides them. Proc. Natl. Acad. Sci. U. S. A. 113, 7650–7654. https://doi.org/10.1073/pnas.1602620113

Geng, Y., Zhao, W., Zhou, F., Ma, X., Yao, S., Hurlemann, R., Becker, B., Kendrick, K.M., 2018. Oxytocin enhancement of emotional empathy: Generalization across cultures and effects on amygdala activity. Front. Neurosci. 12, 512. https://doi.org/10.3389/fnins.2018.00512

Guastella, A.J., Hickie, I.B., McGuinness, M.M., Otis, M., Woods, E.A., Disinger, H.M., Chan, H.K., Chen, T.F., Banati, R.B., 2013. Recommendations for the standardisation of oxytocin nasal administration and guidelines for its reporting in human research. Psychoneuroendocrinology 38, 612–625. https://doi.org/10.1016/j.psyneuen.2012.11.019

Hu, J., Qi, S., Becker, B., Luo, L., Gao, S., Gong, Q., Hurlemann, R., Kendrick, K.M., 2015. Oxytocin selectively facilitates learning with social feedback and increases activity and functional connectivity in emotional memory and reward processing regions. Hum. Brain Mapp. 36, 2132–2146. https://doi.org/10.1002/hbm.22760

Hurlemann, R., Scheele, D., 2016. Dissecting the role of oxytocin in the formation and loss of social relationships. Biol. Psychiatry 79, 185–193. https://doi.org/10.1016/j.biopsych.2015.05.013

Kendrick, K.M., Guastella, A.J., Becker, B., 2017. Overview of human oxytocin research, in: Hurlemann, R., Grinevich, V. (Eds.), Behavioral Pharmacology of Neuropeptides: Oxytocin. Springer, Cham, pp. 321–348. https://doi.org/10.1007/7854_2017_19

Knopp, K., Scott, S., Ritchie, L., Rhoades, G.K., Markman, H.J., Stanley, S.M., 2017. Once a Cheater, Always a Cheater? Serial Infidelity Across Subsequent Relationships. Arch. Sex. Behav. 46, 2301–2311. https://doi.org/10.1007/s10508-017-1018-1

Lansford, J.E., 2009. Parental divorce and children’s adjustment. Perspect. Psychol. Sci. 4, 140–152. https://doi.org/10.1111/j.1745-6924.2009.01114.x

Lee, M.R., Scheidweiler, K.B., Diao, X.X., Akhlaghi, F., Cummins, A., Huestis, M.A., Leggio, L., Averbeck, B.B., 2018. Oxytocin by intranasal and intravenous routes reaches the cerebrospinal fluid in rhesus macaques: determination using a novel oxytocin assay. Mol. Psychiatry 23, 115. https://doi.org/10.1038/mp.2017.27

Li, Q., Becker, B., Wernicke, J., Chen, Y., Zhang, Y., Li, R., Le, J., Kou, J., Zhao, W., Kendrick, K.M., 2019. Foot massage evokes oxytocin release and activation of orbitofrontal cortex and superior temporal sulcus. Psychoneuroendocrinology 101, 193–203. https://doi.org/ https://doi.org/10.1016/j.psyneuen.2018.11.016

Luo, L., Becker, B., Geng, Y., Zhao, Z., Gao, S., Zhao, W., Yao, S., Zheng, X., Ma, X., Gao, Z., Hu, J., Kendrick, K.M., 2017. Sex-dependent neural effect of oxytocin during subliminal processing of negative emotion faces. Neuroimage 162, 127–137. https://doi.org/10.1016/j.neuroimage.2017.08.079

Luo, R., Xu, L., Zhao, W., Ma, X., Xu, X., Kou, J., Gao, Z., Becker, B., Kendrick, K.M., 2017. Oxytocin facilitation of acceptance of social advice is dependent upon the perceived trustworthiness of individual advisors. Psychoneuroendocrinology 83, 1–8. https://doi.org/10.1016/j.psyneuen.2017.05.020

Oliver, B.M., Hyde, S.J., 1993. Gender Differences in Sexuality: A Meta-Analysis. Psychol. Bull. 114, 29–51. https://doi.org/ doi: 10.1037/0033-2909.114.1.29

Paloyelis, Y., Doyle, O.M., Zelaya, F.O., Maltezos, S., Williams, S.C., Fotopoulou, A., Howard, M.A., 2016. A spatiotemporal profile of in vivo cerebral blood flow changes following intranasal oxytocin in humans. Biol. Psychiatry 79, 693–705. https://doi.org/10.1016/j.biopsych.2014.10.005

Penton-Voak, I.S., Perrett, D.I., Castles, D.L., Kobayashi, T., Burt, D.M., Murray, L.K., Minamisawa, R., 1999. Menstrual cycle alters face preference. Nature 399, 741. https://doi.org/10.1038/21557

Place, S.S., Todd, P.M., Penke, L., Asendorpf, J.B., 2010. Humans show mate copying after observing real mate choices. Evol. Hum. Behav. 31, 320–325. https://doi.org/10.1016/j.evolhumbehav.2010.02.001

Preckel, K., Scheele, D., Kendrick, K.M., Maier, W., Hurlemann, R., 2014. Oxytocin facilitates social approach behavior in women. Front. Behav. Neurosci. 8, 191. https://doi.org/10.3389/fnbeh.2014.00191

Quintana, D.S., Westlye, L.T., Alnæs, D.,, Rustan, Ø.G., Kaufmann, T., Smerud, K.T., Mahmoud, R.A., Djupesland, P.G., Andreassen, O.A., 2016. Low dose intranasal oxytocin delivered with Breath Powered device dampens amygdala response to emotional stimuli: A peripheral effect-controlled within-subjects randomized dose-response fMRI trial. Psychoneuroendocrinology 69, 180–188. https://doi.org/10.1016/j.psyneuen.2016.04.010

Quintana, D.S., Westlye, L.T., Smerud, K.T., Mahmoud, R.A., Andreassen, O.A., Djupesland, P.G., 2018. Saliva oxytocin measures do not reflect peripheral plasma concentrations after intranasal oxytocin administration in men. Horm. Behav. 102, 85–92. https://doi.org/10.1016/j.yhbeh.2018.05.004

Scheele, D., Striepens, N., Gunturkun, O., Deutschlander, S., Maier, W., Kendrick, K.M., Hurlemann, R., 2012. Oxytocin modulates social distance between males and females. J. Neurosci. 32, 16074–16079. https://doi.org/10.1523/JNEUROSCI.2755-12.2012

Scheele, D., Striepens, N., Kendrick, K.M., Schwering, C., Noelle, J., Wille, A., Schläpfer, T.E., Maier, W., Hurlemann, R., 2014. Opposing effects of oxytocin on moral judgment in males and females. Hum. Brain Mapp. 35, 6067–6076. https://doi.org/10.1002/hbm.22605

Scheele, D., Wille, A., Kendrick, K.M., Stoffel-Wagner, B., Becker, B., Güntürkün, O., Maier, W., Hurlemann, R., 2013. Oxytocin enhances brain reward system responses in men viewing the face of their female partner. Proc. Natl. Acad. Sci. U. S. A. 110, 20308–20313. https://doi.org/10.1073/pnas.1314190110

Shin, N.Y., Park, H.Y., Jung, W.H., Kwon, J.S., 2018. Effects of Intranasal Oxytocin on Emotion Recognition in Korean Male: A Dose-Response Study. Psychiatry Investig. 15, 710. https://doi.org/10.30773/pi.2018.02.19

Spengler, F.B., Schultz, J., Scheele, D., Essel, M., Maier, W., Heinrichs, M., Hurlemann, R., 2017. Kinetics and dose dependency of intranasal oxytocin effects on amygdala reactivity. Biol. Psychiatry 82, 885–894. https://doi.org/10.1016/j.biopsych.2017.04.015

Striepens, N., Kendrick, K.M., Hanking, V., Landgraf, R., Wüllner, U., Maier, W., Hurlemann, R., 2013. Elevated cerebrospinal fluid and blood concentrations of oxytocin following its intranasal administration in humans. Sci. Rep. 3, 3440. https://doi.org/10.1038/srep03440

Takahashi, H., Matsuura, M., Yahata, N., Koeda, M., Suhara, T., Okubo, Y., 2006. Men and women show distinct brain activations during imagery of sexual and emotional infidelity. Neuroimage 32, 1299–1307. https://doi.org/10.1016/j.neuroimage.2006.05.049

Unkelbach, C., Guastella, A.J., Forgas, J.P., 2008. Oxytocin selectively facilitates recognition of positive sex and relationship words. Psychol. Sci. 19, 1092–1094. https://doi.org/10.1111/j.1467-9280.2008.02206.x

Xu, L., Ma, X., Zhao, W., Luo, L., Yao, S., Kendrick, K.M., 2015. Oxytocin enhances attentional bias for neutral and positive expression faces in individuals with higher autistic traits. Psychoneuroendocrinology 62, 352–358. https://doi.org/10.1016/j.psyneuen.2015.09.002

Zhao, W., Geng, Y., Luo, L., Zhao, Z., Ma, X., Xu, L., Yao, S., Kendrick, K.M., 2017. Oxytocin increases the perceived value of both self-and other-owned items and alters medial prefrontal cortex activity in an endowment task. Front. Hum. Neurosci. 11, 272. https://doi.org/10.3389/fnhum.2017.00272

Zhao, W., Ma, X., Le, J., Ling, A., Xin, F., Kou, J., Zhang, Y., Luo, R., Becker, B., Kendrick, K.M., 2018. Oxytocin biases men to be more or less tolerant of others’ dislike dependent upon their relationship status. Psychoneuroendocrinology 88, 167–172. https://doi.org/10.1016/j.psyneuen.2017.12.010

## References

Baron-Cohen, S., Wheelwright, S., Skinner, R., Martin, J., Clubley, E., 2001. The autism-spectrum quotient (AQ): Evidence from asperger syndrome/high-functioning autism, malesand females, scientists and mathematicians. J. Autism Dev. Disord. 31, 5–17. https://doi.org/10.1023/A:1005653411

Beck, A.T., Steer, R.A., Brown, G.K., 1996. Beck depression inventory-II. The Psychological Corporation, San Antonio, TX.

Berry, J.W., Worthington, E.L., O’Connor, L.E., Parrott, L., Wade, N.G., 2005. Forgivingness, vengeful rumination, and affective traits. J. Pers. 73, 183–226. https://doi.org/10.1111/j.1467-6494.2004.00308.x

Brown, R.P., 2003. Measuring Individual Differences in the Tendency to Forgive: Construct Validity and Links with Depression Measuring Individual Differences in the Tendency to Forgive: Construct Validity and Links With Depression. Personal. Soc. Psychol. Bull. 29, 759–771. https://doi.org/10.1177/0146167203029006008

Costa, P., Mccrae, R.R., 1989. The NEO-PI/NEO-FFI manual supplement. Psychological Assessment Resources, Odessa, FL.

Davis, M.H., 1980. A multidimensional approach to individual differences in empahty. Cat. Sel. Doc. Psychol. 40, 3480.

Hatfiled, E., Sprecher, S., 1986. Measuring passionate love in intimate relations. J. Adolesc. 9, 383–410. https://doi.org/10.1016/S0140-1971(86)80043-4

Hendrick, C., Hendrick, S., 1986. A theory and method of love. J. Pers. Soc. Psychol. 50, 392–402. https://doi.org/10.1037/0022-3514.50.2.392

Liebowitz, M.R., 1987. Social phobia, in: Anxiety. Karger Publishers, pp. 141–173. https://doi.org/10.1159/000414022

Rosenberg, M., 1965. Society and the adolescent self-image. Princeton university press Princeton, NJ.

Siegrist, M., Keller, C., Barle, T.C., Gutscher, H., 2005. Effects of general trust on cooperation in the investment game and in a social dilemma Unpublished manuscript, Institute for Environmenta.

Watson, D., Clark, L.A., Tellegen, A., 1988. Development and validation of brief measures of positive and negative affect: the PANAS scales. J. Pers. Soc. Psychol. 54, 1063–1070. https://doi.org/10.1037/0022-3514.54.6.1063

